# Homology Modeling and Docking Study of Shewanella Like Protein Phosphatase Involved in Development of Ookinetes in Plasmodium

**DOI:** 10.1101/297770

**Authors:** Sandhini Singh, Ruchi Yadav

## Abstract

Parasites of the genus Plasmodium cause a great deal of morbidity and mortality in worldwide, largely in regions with limited access and indication to the tools necessary to control mosquito populations and to treat human infections of Malaria. Five species of this class of eukaryotic pathogens cause different human disease, with Plasmodium falciparum alone is infecting approximately 500 million people per year and resulting in approximately 1 million deaths. The two genes encoding the Shewanella-like phosphatases of *P.falciparum*, SHLP-1 and SHLP-2, are conserved among members of Plasmodium family. SHLP is frequently found in asexual blood stages and expressed at all stages of the life cycle of parasite. SHLP deletion results in a reduction in microneme formation, ookinetes (zygote) development and complete ablation of oocyst formation, thereby blocking transmission of parasite.. Structure Modeling of SHLP protein can be helpful in understanding the active site and binding site information and hence can be used for drug designing for therapeutics against malaria. Effective role of Resveratrol is studied against SHLP protein using Docking method to identify protein-ligand interaction scheme and bond formation. Study suggests that resveratrol have strong interaction with SHLP protein and can be used as ligand for drug designing.

## 1. INTRODUCTION

The unicellular parasites of the genus *Plasmodium* are the causative agent of malaria, and results shows with annual fatalities close to 1.25 million. Malaria is caused by infection with the apicomplexan parasite *Plasmodium*, is transmitted via the female *Anopheles* mosquito and in 2012 resulted in approximately 207 million clinical infections and over 600,000 deaths [1]. The life cycle of *Plasmodium* progresses through different morphologically distinct stages of development, including asexual proliferation in hepatocytes, followed by clinically overt intra erythrocytic multiplication in the vertebrate host [2]. Ingestion of developmentally arrested gametocytes triggers development of sexual the parasite in the mosquito, with eventual migration to the salivary glands and transmission during process of feeding [3]. During each stage the parasite utilizes and control a number of signal transduction mechanisms, including reversible protein phosphorylation catalyzed by protein kinases (PKs) and phosphatases (PPs). This signaling mechanism is a conserved, ubiquitous regulatory process for many prokaryotic and eukaryotic cellular pathways [4]. However, while PKs are well recognized as important therapeutic targets, PPs are only now emerging as targets for the purpose of clinical intervention [5].

### Shewanella-like Protein Phosphatase (SHLP)

Mammalian *Plasmodium* development process initiate species proceeds via asexual exoerythrocytic proliferation and intraerythrocytic multiplication that existing in mammalian liver hepatocytes and erythrocytes, respectively, whereas sporogony and sexual development take place in the mosquito [6]. *Plasmodium* is related to the phylum *Apicomplexa*, which is characterized and indicated by the distinct apical organelles presence consisting of micronemes, dense granules, and rhoptries that are used by the parasite for gliding motility and host invasion[7] Of the three invasive stages (sporozoites, merozoites, and ookinetes), the ookinetes uniquely lacks rhoptries and dense granules[8]. There are two nonconventional PPs, one of them is containing an N-terminal β-propeller formed by kelch-like motifs and the other a *Shewanella-*like PP (SHLP1), are required during ookinetes-to-oocyst transition and subsequent transmission of the malaria through *P. berghei* [9]. More recently, discussed a second SHLP member of family (SHLP2) was implicated in dephosphorylation of the host protein Band 3 during merozoites invasion of erythrocytes [10]. In eukaryotes other than in *Apicomplexa*, SHLPs have been found in, *Archaeplastida*, some fungi, and some *Chromalveolates* and are structurally related to a segment of bacterial PPPs those are identified firstly in the psychrophilic bacteria *Shewanella* and *Colwellia* [11]. the two genes presence in the *P. berghei* genome resulted encoding *Shewanella*-like PPs was confirmed in first chance, which has a signal peptide and apicoplast targeting sequence and second, which were named *Shewanella*-like protein phosphatases SHLP1 and SHLP2[12]. The analysis of *Plasmodium* SHLPs explained that only the highly conserved STP catalytic residues and binding sites of metal ion are largely conserved [13]. The rest of the conserved STP residues, okadaic acid, and microcystin inhibitor binding sites, as well as PP1 regulatory subunit binding sites and PP1 substrate binding sites are only partially or not conserved[14]. SHLP1 plays an important (though not essential) role at an early stage in ookinetes development and differentiation [15]. SHLP1 in *Plasmodium* is essential for parasite transmission via the mosquito and thus a potential target for the development of transmission-blocking drugs. Ookinetes-to-oocyst transition represents one of the biggest bottlenecks in the life cycle of the malarial parasite, and SHLP1 plays a crucial role in this current process [16].

### Potential Inhibitors of SHLP protein

Reversible protein phosphorylation is of central importance to the proper cellular functioning of all living organisms [17]. Catalyzed by the opposing reactions of protein kinases and phosphatases, dysfunction in reversible protein phosphorylation can result in a wide variety of cellular aberrations [18].

An antioxidant is a molecule that inhibits the oxidation of other molecules [19]. The term "antioxidant" is mainly used for two different groups of substances: chemical industry which are perused to products to prevent oxidation, and natural chemicals found in foods and body tissue which are said to have beneficial health effects [20]. Resveratrol, a natural product, is known to affect a broad range of intracellular mediators and also possess some anti-oxidant activity [21]. Resveratrol (3,5,4′-trihydroxy-*trans*-stilbene) is a stilbenoid, a type of natural phenol, and a phytoalexin produced by several plants in response to injury or when the plant is under attack by pathogens such as bacteria or fungi. Sources of resveratrol in food include the skin of grapes, blueberries, raspberries, mulberries, lingonberry and senna [22]. Resveratrol is produced in plants by the action of the enzyme, resveratrol synthase [23].

## 2. MATERIALS AND METHOD

Study of SHLP and its variants is done using UniprotKB database (http://www.uniprot.org/). Homology Modeling was done using Schrödinger software and structure verified using Ramachandran plot. Ten antioxidants were searched in Pubchem database (https://pubchem.ncbi.nlm.nih.gov/). All modeled proteins were docked with ten ligands using Glide Docking program of Schrödinger software suite. Docking results are analyzed and protein –ligand interaction map is studied to identify best ligand and interaction energy.

### Target Protein

SHLP protein is searched in UniProt database and two types of SHLP protein were found SHLP1 and SHLP2. Table 1 list details of SHLP protein, UniProt ID, Gene name and source organism.

**Table 1.** SHLP Protein identified form UniProt database.

| S.No. | Uniprot ID | Gene Name | Organism |
| --- | --- | --- | --- |
| 1. | A0A1C3KEC2 | SHLP 2 | *Plasmodium sp.* |
| 2. | A0A1D3TM00 | SHLP 2 | *Plasmodium ovale (malaria parasite P. ovale)* |
| 3. | A0A060RV45 | SHLP 2 | *Plasmodium reichenowi* |
| 4. | A0A1C6X2Z2 | SHLP1 | *Plasmodium berghei* |

### Ligands for Docking

Ligands were retrieved from Pubchem database for interaction analysis with modeled SHLP protein.Table 3 list the details of antioxidants ligand used for docking all four homology modeled protein (modeled protein A, B, C, D).

**Table 2.** List of Antioxidants used for Docking from Pubchem database.

| S.No . | Pubchem CID | Chemical Name | Molecular Formula | Molecular Weight | Log P Value | H-bond Donor | H-bond Acceptor |
| --- | --- | --- | --- | --- | --- | --- | --- |
| 1. | 637098 | Maximol A | C<sub>28</sub>H<sub>22</sub>O<sub>6</sub> | 454.478 g/mol | 5.4 | 5 | 6 |
| 2. | 71752006 | Trans Resveratrol Penta-O-Acetyl-3-β-D-Glucuronide Methyl Ester | C<sub>31</sub>H<sub>32</sub>O<sub>14</sub> | 628.583 g/mol | 3.3 | 0 | 14 |
| 3. | 5473050 | Pinostilbene | C<sub>15</sub>H<sub>14</sub>O<sub>3</sub> | 242.274 g/mol | 3.5 | 2 | 3 |
| 4. | 75055173 | Resveratrol phosphonate | C<sub>14</sub>H<sub>12</sub>Na<sub>3</sub>O<sub>12</sub>P<sub>3</sub> | 534.129 g/mol | N/A | 3 | 12 |
| 5. | 445154 | Resveratrol | C<sub>14</sub>H<sub>12</sub>O<sub>3</sub> | 228.247 g/mol | 3.1 | 3 | 3 |
| 6. | 53394008 | Resveratrol 3-O-D-Glucuronide | C<sub>20</sub>H<sub>20</sub>O<sub>9</sub> | 404.371 g/mol | 1.6 | 6 | 9 |
| 7. | 101160946 | Resveratrol Cis-Dehydrodimer | C<sub>28</sub>H<sub>22</sub>O<sub>6</sub> | 454.478 g/mol | 5.4 | 5 | 6 |
| 8. | 86664995 | Lycopene Resveratrol | C<sub>54</sub>H<sub>68</sub>O<sub>3</sub> | 765.135 g/mol | N/A | 3 | 3 |
| 9. | 86662966 | Resveratrol Tripalmitate | C<sub>62</sub>H<sub>108</sub>O<sub>9</sub> | 997.537 g/mol | N/A | 6 | 9 |
| 10. | 46855201 | Resveratrol<br>Potassium4,-<br>Sulfate | $C_{14}H_{11}KO_6$<br>S | 346.394<br>g/mol | N/A | 2 | 6 |

**Table 3.**
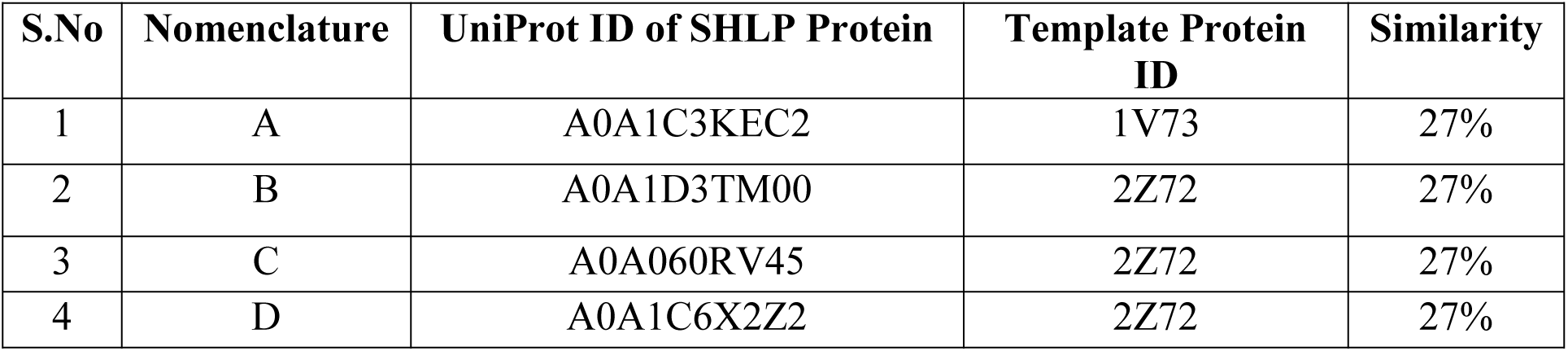
Template details of SHLP protein.

## 3. RESULTS AND DISCUSSION

**1) Homology Modeling using Schrödinger software**

*Shewanella*-like protein phosphatase (SHLP1) is an ancient bacterial protein also found in fungi, protists, and plants.SHLP1 protein in malarial parasite plasmodium is involved in ookinetes (zygote) development, microneme formation and it is found abundantly in asexual stages of parasitic life cycle. Here we predict structure of SHLP protein by homology modeling and structure conservation in other variants of SHLP protein found in different species of plasmodium.

Homology Modeling of all 4 proteins was done and modeled was obtained using Schrodinger Software Suite. The modeled structures and their respective Ramachandran Plot are specified in the figures (1-8).

### Modeled structure of protein A(SHLP 2, UniProt ID: A0A1C3KEC2)

**Figure 1.**
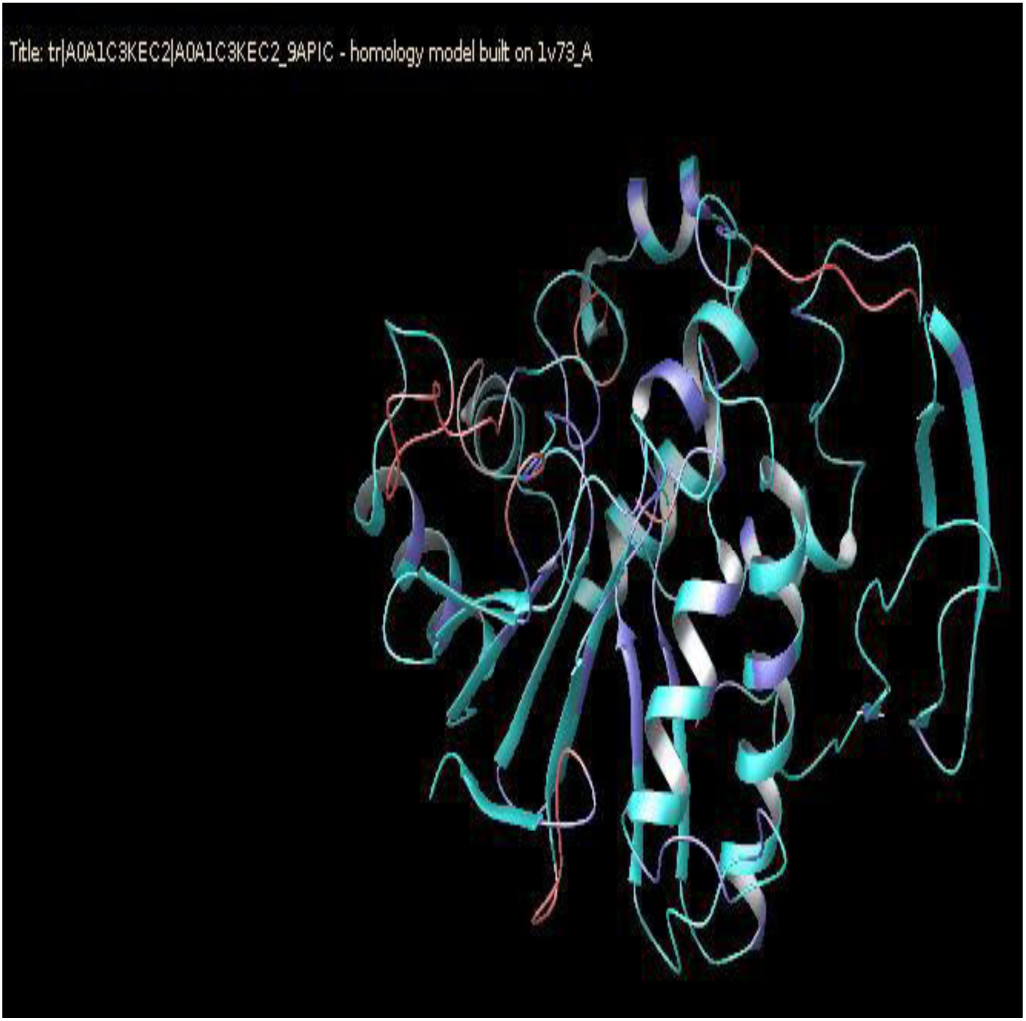
Result of Schrödinger Maestro showingthe showing the Ramachandran Plot for protein A.

**Figure 2.**
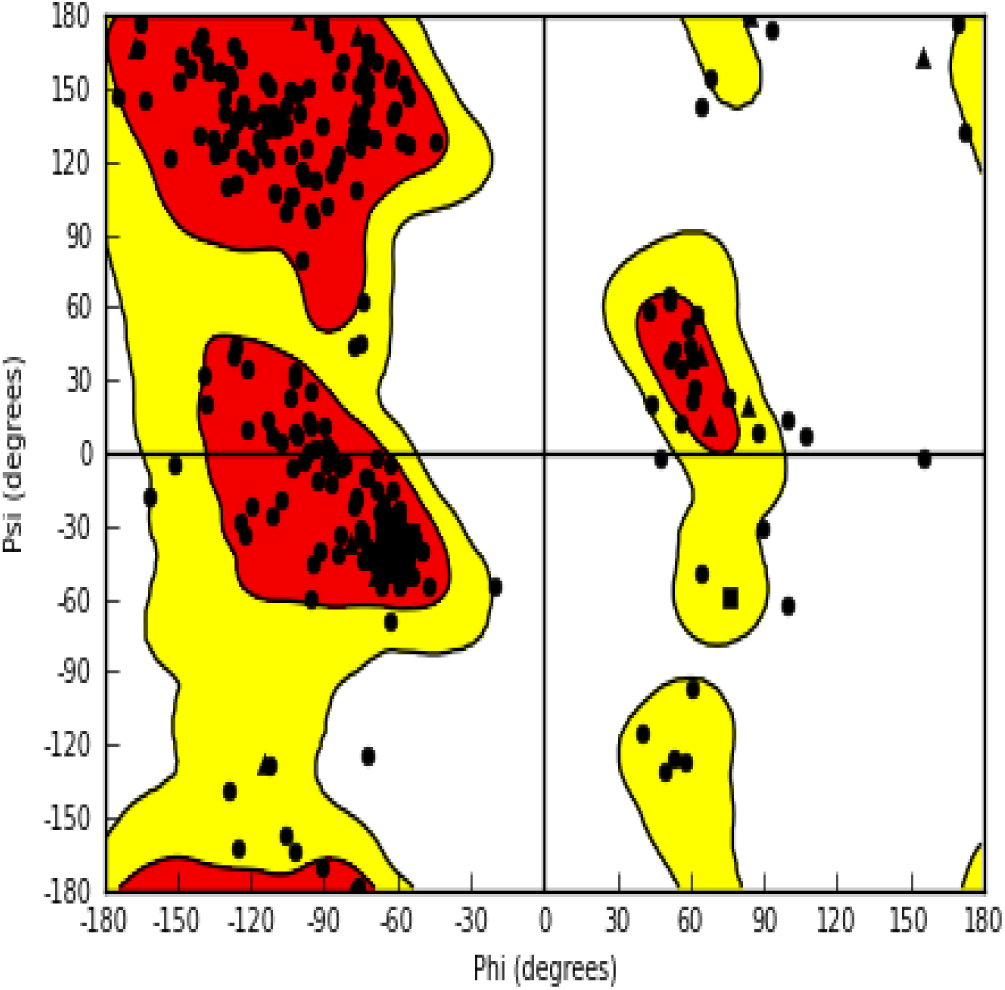
Result of Schrödinger Maestro in-silico modeled structure of protein A.

### Modeled structure of protein B(SHLP 2, UniProt ID: A0A1D3TM00)

**Figure 3.**
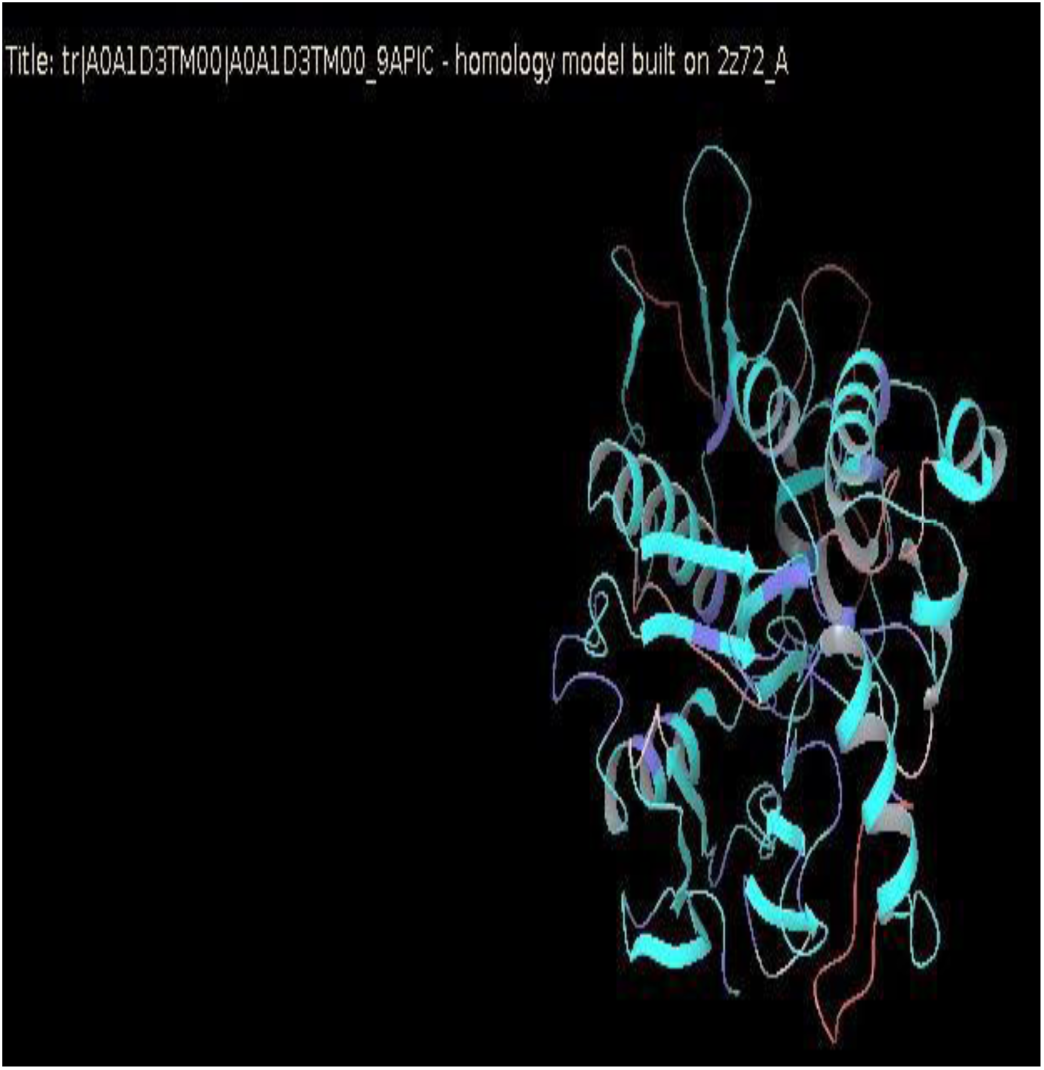
Result of Schrödinger Maestro showing the in-silico showing the Ramachandran Plot for protein B.

**Figure 4.**
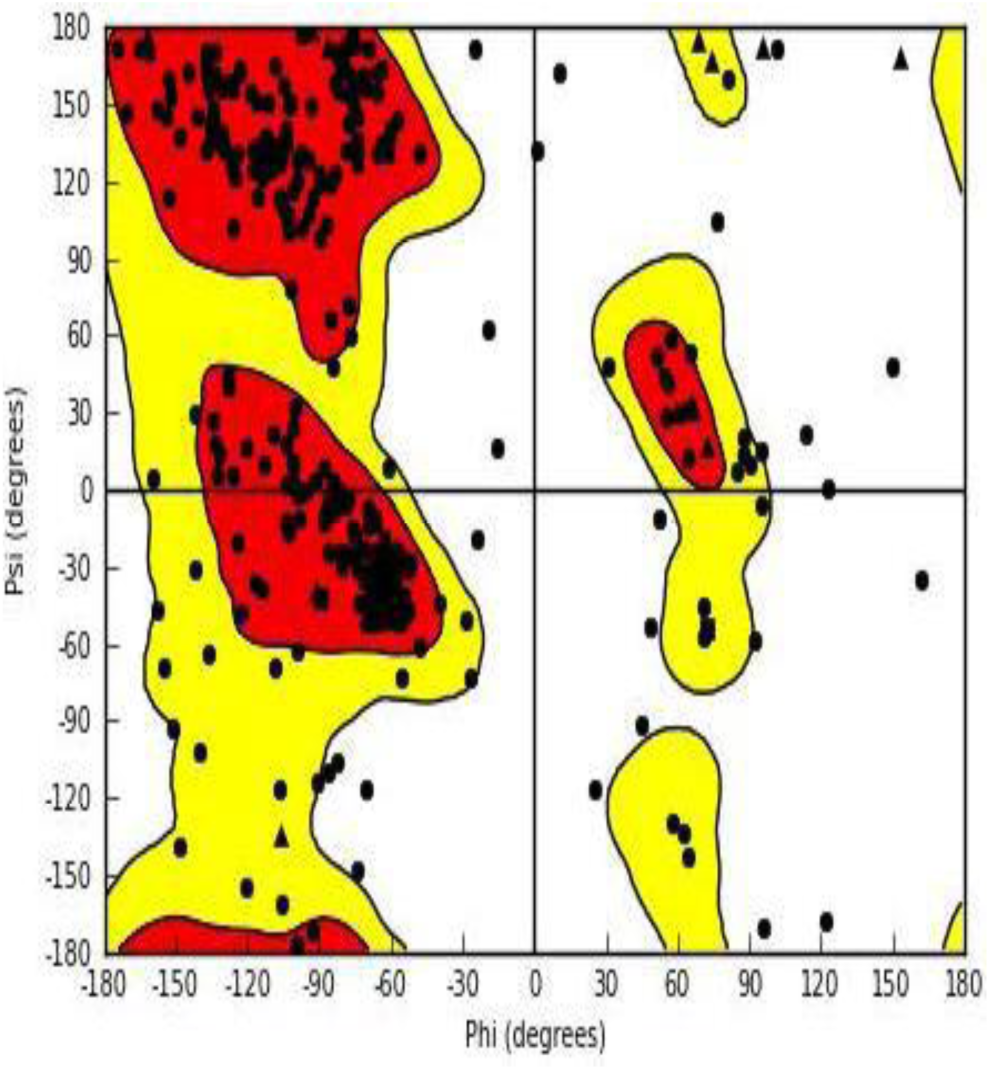
Result of Schrödinger Maestro modeled structure of protein B.

### Modeled structure of protein C (SHLP 2, UniProt ID: A0A060RV45)

**Figure 5.**
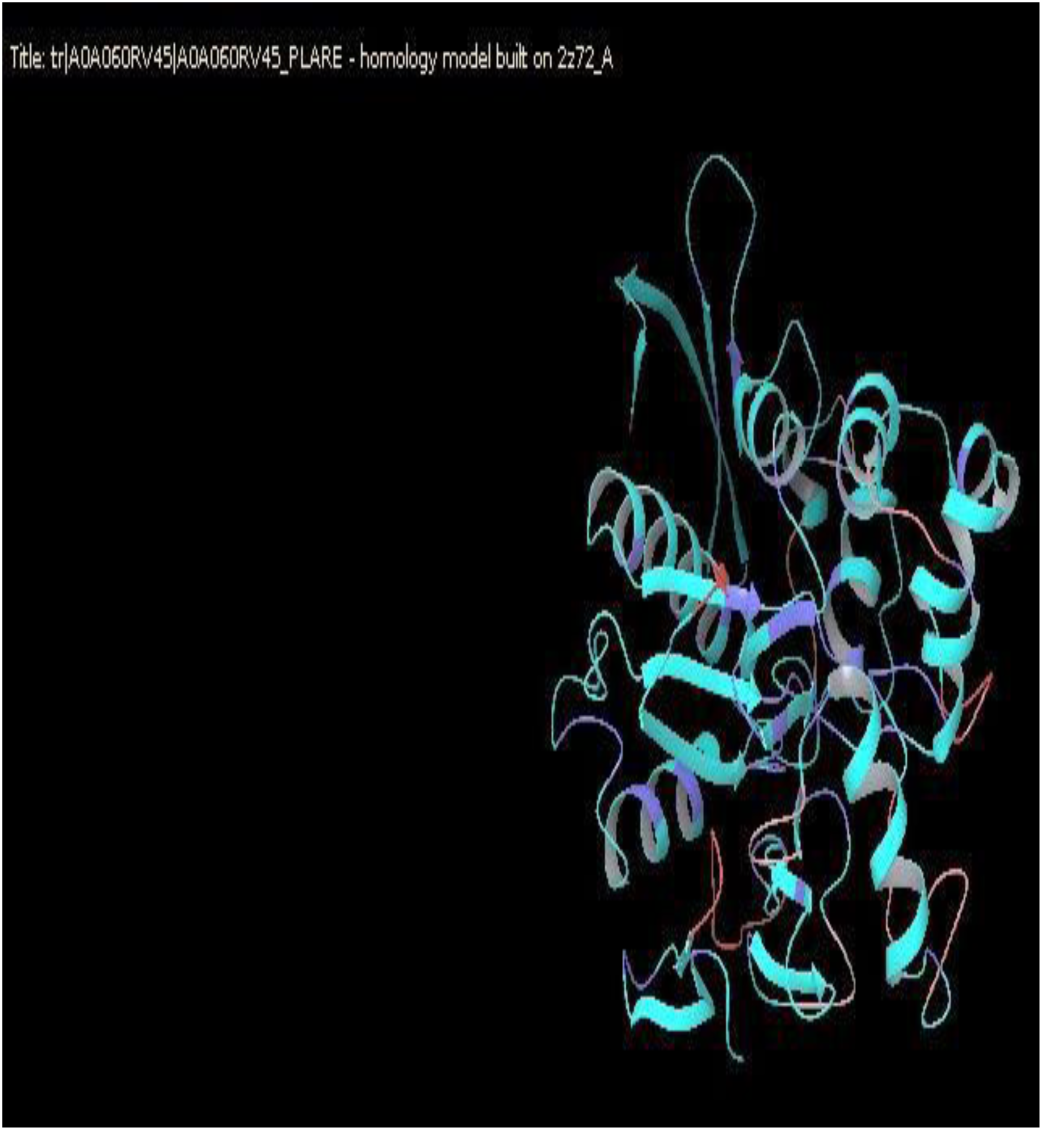
Result of Schrödinger Maestro showing the in-silico Ramachandran Plot for protein C.

**Figure 6.**
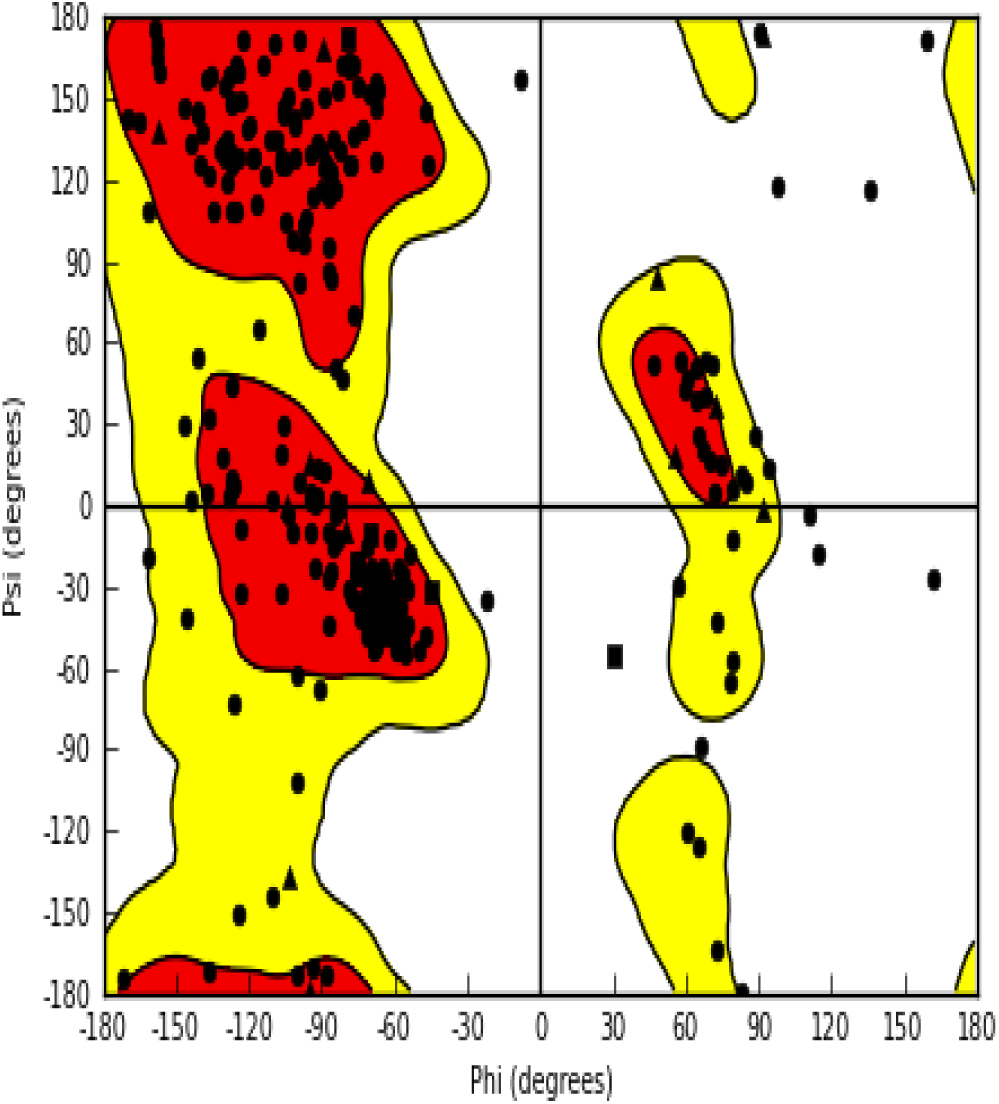
Result of Schrödinger Maestro showing the modeled structure of protein C.

### Modeled structure of protein D (SHLP 1, UniProt ID: A0A1C6X2Z2)

**Figure 7.**
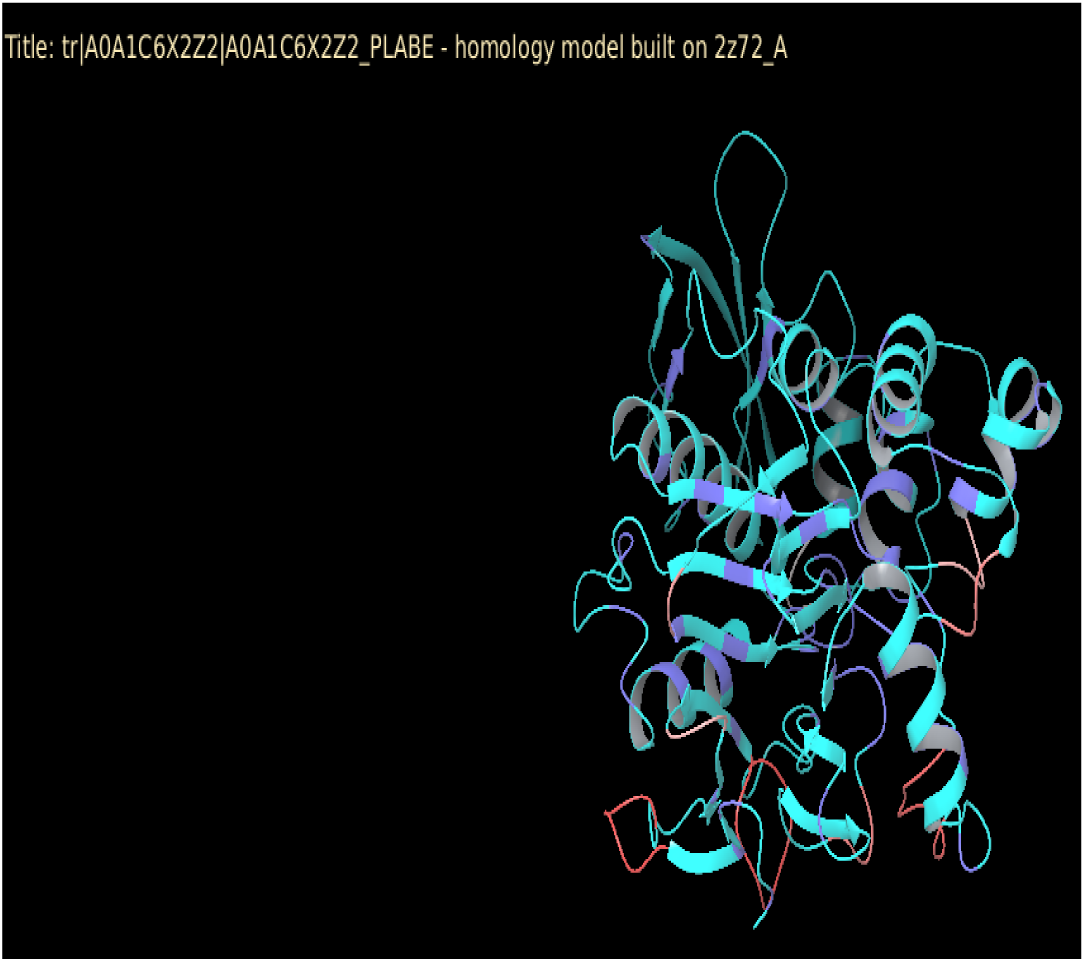
Result of Schrödinger Maestro showing the in-silico Ramachandran Plot for protein D.

**Figure 8.**
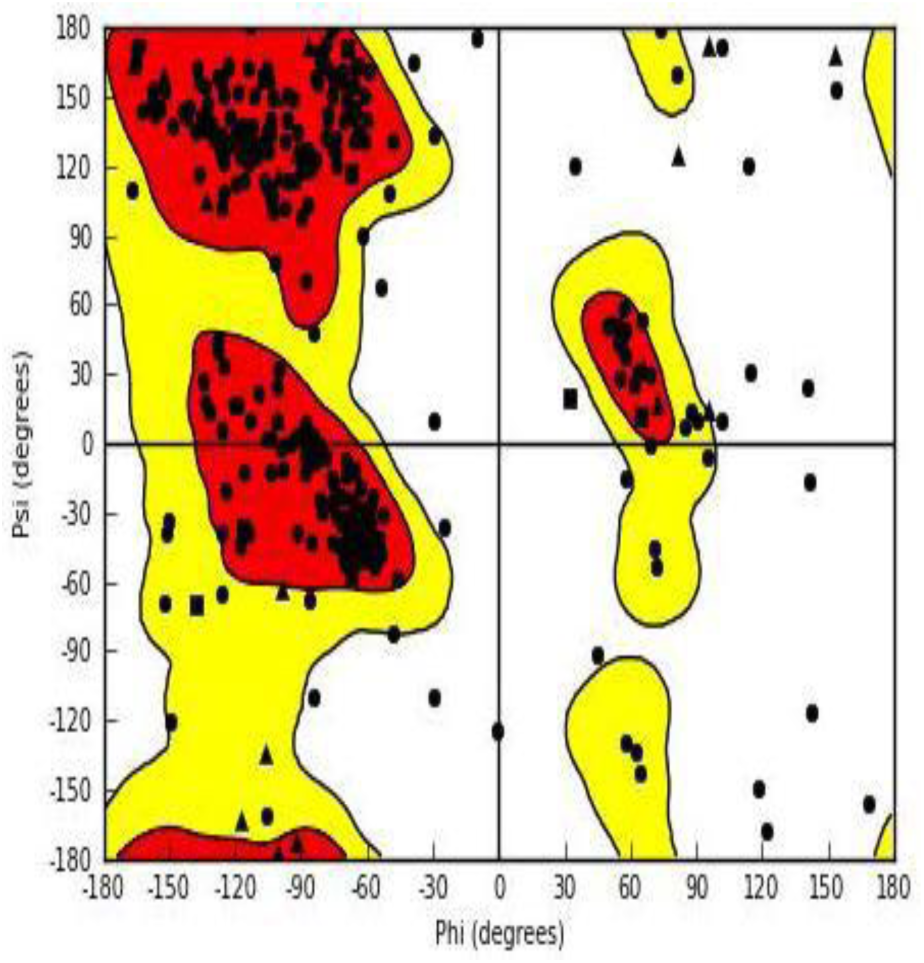
Result of Schrödinger Maestro showing the modeled structure of protein D.

Two variants of SHLP protein (SHLP1 and SHLP2) was found in four different species of plasmodium *(Plasmodium sp., Plasmodium ovale, Plasmodium reichenowi, and Plasmodium berghei*) retrieved through UniProt database. Homology modeling of SHLP1 and SHLP2 protein was done using Schrödinger software and phyre server. Four template viz: 1V73, 2Z72, 2Z72, 2Z72 is used to model structure of SHLP protein (Fig 1-8) and structure verification is done using Ramachandran plot shows approximately 86% residues in favoured region for all four SHLP protein.. This structure is used for docking studies to identify potential inhibitor against SHLP protein.

## 3. Docking

Ligand retrieved from Pubchem database for docking was Resveratrol. Resveratrol is a polyphenolic compound which has been derived from plants. Resveratrol activates sirtuin1 (an enzyme which deacetylates the proteins which contribute to the regulation of cells) which further binds to PGC-1 alpha (a transcriptional coactivator which takes care of biogenesis and functioning of mitochondria) and activates it by the process of deacetylation

Docking result shows that resveratrol have a strong binding to SHLP protein. Ligand protein interaction map as shown in table 4 shows that resveratrol make hydrogen bonds with SHLP protein. Hence resveratrol can be potential drug against Malaria caused by development and proliferation of ookinetes in *Plasmodium*.

**Table 4.** Docking result of SHLP protein with Ligands and Ligand-protein interaction map.

| S.NO | Protein | Best Ligand | Glide Score | Bond Formation | Ligand-Protein Interaction Map |
| --- | --- | --- | --- | --- | --- |
| 1. | SHLP 2<br>(A0A1C3KEC2) | Pinostilbene | -6.80 | Leu 72,<br>Hydrogen |  |
| 2. | SHLP 2<br>(A0A1D3TM00) | Resveratrol<br>Tripalmitate | -0.72 | Tyr 315,<br>Ester |  |
| 3. | SHLP 2<br>(A0A060RV45) | Resveratrol<br>Tripalmitate | -<br>0.308 | Asn 127,<br>Tyr 130,<br>Gly 131,<br>Ester |  |
| 4. | SHLP1<br>(A0A1C6X2Z2) | Resveratrol<br>Tripalmitate | -1.24 | Lys 235,<br>Ester |  |

## 4. CONCLUSION

In most organisms, differentiation, and development processes involved in cell cycle, are regulated by proteins of reversible phosphorylation. While kinases are well identified as basic targets for drugs, the SHLPs study reports has only recently begun to identify them as therapeutic targets of potential activity. Variants of SHLP from different *Plasmodium* species were studied and their structure was modeled using Schrödinger. SHLP plays a crucial role in ookinetes and microneme development in *Plasmodium* so ligand which can interact and inhibit SHLP protein, can be potential drug against malarial parasite development. We studied the binding of anti-oxidant, like Resveratrol, with this protein using docking method and it was found that resveratrol an anti-oxidant can bind with the target SHLP protein. This study can be helpful in designing drugs against SHLP protein hence can be used as drug against malaria.

## 5. ACKNOWLEDGEMENT

I acknowledge to all those who are directly or indirectly helped me in this work. Further I wish to acknowledge Bioinformatics analysis tools and data bases for conducting this study.

